# Autism-associated *ASPM* variant causes macrocephaly and social-cognitive deficits in mice

**DOI:** 10.1101/2025.02.17.638753

**Authors:** Sonu Singh, Hyopil Kim, Alev Ecevitoglu, Renee Chasse, Alexis M. Ludko, Basavaraju Sanganahalli, Vishaak Gangasandra, Sung Rye Park, Siu-Pok Yee, James Grady, John Salamone, R. Holly Fitch, Timothy Spellman, Fahmeed Hyder, Byoung-Il Bae

## Abstract

In autism spectrum disorder (ASD), a neurodevelopmental disorder with social-cognitive deficits, macrocephaly occurs in 20% of patients with severe symptoms. However, the role of macrocephaly in ASD pathogenesis remains unclear. Here, we address the mechanistic link between macrocephaly and ASD by investigating a novel ASD-associated gain-of-function A1877T mutation in *ASPM* (*abnormal spindle-like microcephaly-associated*). *ASPM* is a key regulator of cortical size and cell proliferation expressed in both excitatory and inhibitory neuronal progenitors but not in differentiated neurons. We found that *Aspm* gain-of-function knock-in mice exhibit macrocephaly, excessive embryonic neurogenesis with expanded outer radial glia, an increased excitatory-inhibitory (E-I) ratio, brain hyperconnectivity, and social-cognitive deficits with male specificity. Our results suggest that macrocephaly in ASD is not a proportional expansion of excitatory and inhibitory neurons, but a shift in the E-I ratio, independent of the expression patterns of the causative gene. Thus, macrocephaly alone can cause a subset of ASD-like symptoms.

## INTRODUCTION

Autism spectrum disorder (ASD) is a genetic neurodevelopmental disorder with social-cognitive deficits and distinct comorbidities^1^. One such comorbidity is macrocephaly (“large head”), a head circumference exceeding the mean by >2 standard deviations, observed in approximately 20% of patients exhibiting severe ASD symptoms, including intellectual disability^2-7^. Postmortem examination of the prefrontal cortex in affected children reveals a 67% increase in neuron number relative to controls, while neuron size remains unchanged^3^. Similarly, patient-derived cortical organoids display an increase in size^8, 9^. Notably, both head circumference and cortical organoid size positively correlate with ASD symptom severity. However, the mechanistic relationship between macrocephaly and ASD remains unresolved.

Macrocephaly results from dysregulated expansion of the cortical progenitor pool, which ultimately determines neuronal number. Mouse models harboring haploinsufficiency in the ASD risk genes *Pten* and *Chd8* exhibit increased progenitor proliferation and macrocephaly^10, 11^. However, as these genes are expressed in both cortical progenitors and postmitotic neurons, the extent to which macrocephaly itself contributes directly to ASD symptoms is unclear.

The *ASPM* (*abnormal spindle-like microcephaly associated*) gene is a primary determinant of cerebral cortical size, with early-truncating loss-of-function (LOF) mutations being a leading cause of genetic microcephaly (“small head”)^12, 13^. *ASPM* encodes a centrosomal protein that is expressed exclusively in proliferating cells, including cortical progenitors, but not in terminally differentiated, postmitotic neurons. Loss of *ASPM* reduces cortical progenitor proliferation in multiple species, including zebrafish^14^, mice^15-21^, ferrets^22, 23^, and human cortical organoids^24-26^. Conversely, *ASPM* overexpression is associated with increased cell proliferation and implicated in cancer^27, 28^, consistent with a report that genetic variants linked to head size share common pathways with cancer^29^. Based on these observations, we hypothesize that gain-of-function (GOF) mutations in *ASPM*, which are associated with both ASD and cancer, may drive macrocephaly by enhancing progenitor cell proliferation without directly affecting neurons. Identifying such variants could clarify the specific role of macrocephaly in ASD.

Analysis of ASD and cancer gene databases^30, 31^ reveals a few missense variants in *ASPM* that are associated with both conditions. Among these, we found that the A1877T mutation^32^, a substitution of alanine with threonine at position 1877, significantly upregulates ASPM protein levels, suggesting a GOF effect. To investigate further, we generated knock-in (KI) mice carrying the orthologous mutation. The KI mice exhibit macrocephaly, expanded cortical progenitor pool including increased outer radial glia (oRG)-like cells, increased excitatory-inhibitory (E-I) ratio, brain hyperconnectivity, and social-cognitive deficits. These findings suggest that macrocephaly alone may be sufficient to account for a subset of ASD symptoms.

## RESULTS

### Identification of an *ASPM* GOF variant associated with ASD and generation of KI mice

Identifying a GOF variant of *ASPM* associated with ASD may elucidate the role of macrocephaly in ASD. To identify a potential *ASPM* GOF variant, we first analyzed the SFARI database of ASD-associated genes^30^ and identified 28 missense variants reported in individuals with ASD (Supplementary Table 1). Given that ASPM is a rapidly evolving protein^33^, we refined our selection to 18 variants in which the wild-type (WT) amino acids are conserved across vertebrates, indicating functional significance.

Since *ASPM* regulates cell proliferation, a GOF variant could also be linked to cancer. To investigate this possibility, we cross-referenced these variants with the COSMIC cancer gene database^31^ and identified four—R942C, R1667H, A1877T, and A1950V—each reported in at least two cancer cases. Notably, these variants are located either within the calponin homology domain or clustered within the IQ calmodulin-binding domain repeats, both essential for ASPM function. Among these, A1877T (c.5629G>A, ClinVar VCV000157838.18) has been reported in seven cases of ASD^32^ and three cases of lung cancer^34^, making it a strong GOF candidate for further investigation (Fig. 1a-1b).

**Figure 1.**
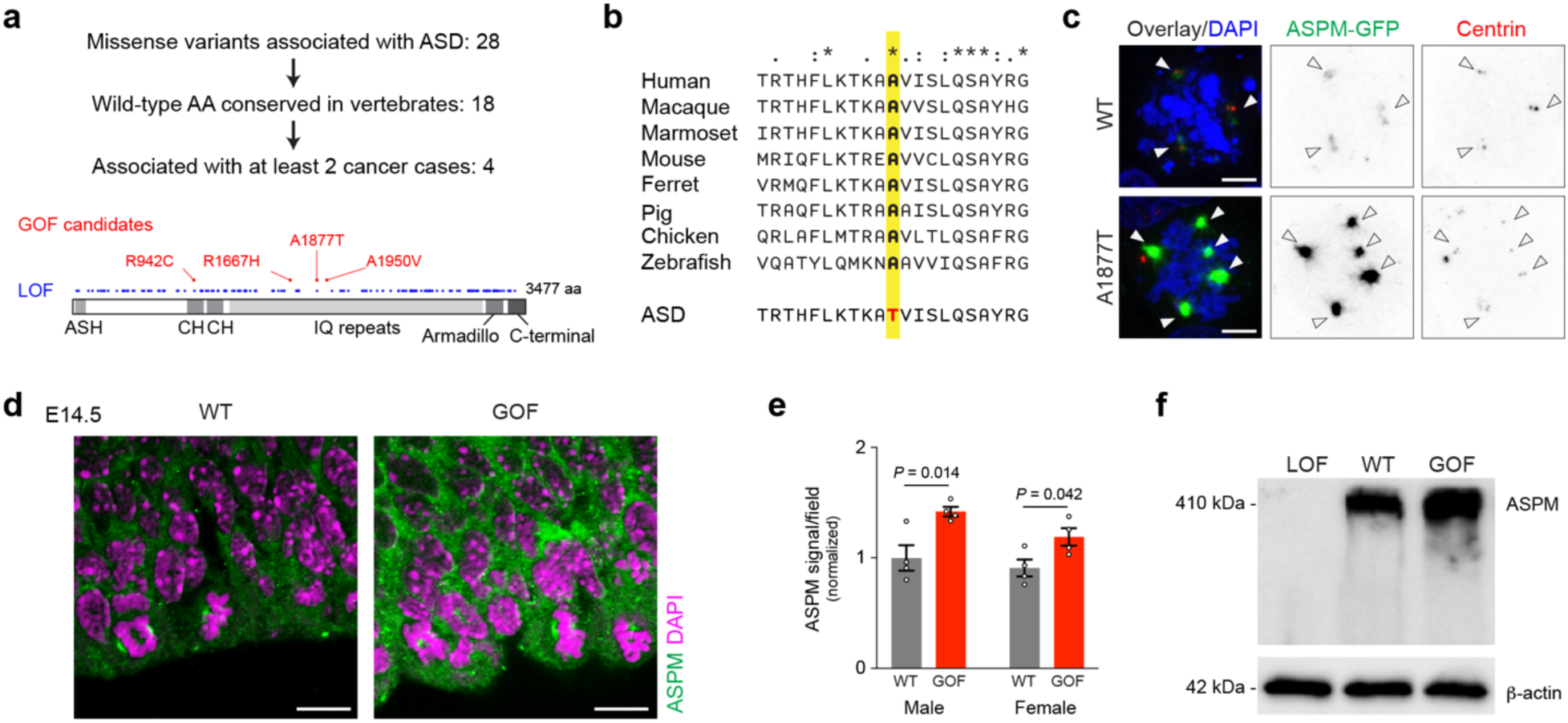
Identification of an ASD-associated GOF variant in *ASPM* and generation of KI mice. **(a)** Selection of four potential GOF missense variants in *ASPM*. These variants are localized within either the CH domain or IQ domain repeats. In contrast, early-truncating LOF mutations are distributed throughout the protein^54^. **(b)** Alignment of the ASPM protein across humans and seven model organisms reveals strong conservation of Ala1877, the residue mutated in ASD. **(c)** Exogenous expression of ASPM-GFP in 293T cells, without or with the A1877T mutation, demonstrates a robust GOF effect while preserving centrosomal localization. Scale bar, 5 μm. **(d-f)** Immunostaining and Western blot analysis in homozygous Aspm GOF KI mice at E14.5 show a 1.5-fold increase in ASPM protein levels compared to WT. N = 4/genotype. Student’s *t*-test. Error bars, mean ± s.e.m. Scale bars, 10 μm.

To assess the effect of A1877T on ASPM protein, we generated the mutant construct and compared the expression levels of WT and A1877T ASPM-GFP in heterologous cells. The A1877T variant exhibits significantly increased expression levels compared to WT while maintaining centrosomal localization (Fig. 1c), supporting its classification as a robust *ASPM* GOF variant associated with ASD and a potential driver of macrocephaly.

To develop a model system to investigate the *in vivo* effects of the *ASPM* GOF variant, we generated a KI mouse model on a C57BL/6 background carrying the orthologous A1845T mutation using genome editing (see Methods). Both heterozygous and homozygous KI mice, of both sexes, are viable and fertile. Immunostaining and Western blot analysis of embryonic day 14.5 (E14.5) homozygous KI mouse brains reveal a 1.5-fold increase in ASPM protein levels, confirming the GOF effect *in vivo* (Figs. 1d–1f). Although the GOF variant is expected to exert a dominant effect over the WT allele, and the seven reported ASD patients carrying this variant are heterozygous^32^, we opted to study homozygous KI mice to obtain more robust phenotypes. Thus, we have established a novel mouse model carrying an ASD-associated GOF variant of *Aspm*.

### *Aspm* GOF mice show macrocephaly with increased neuronal density

Whether the *ASPM* GOF variant causes macrocephaly was the first question we addressed. We found that, in both male and female GOF mice, forebrain weight is significantly increased at postnatal days 1 (P1) and 10 (P10), while the overall body weight remains unchanged (Figs. 2a–2c). However, by P50, these differences in forebrain weight are no longer significant. To more precisely assess cortical size, we performed non-invasive structural magnetic resonance imaging (MRI) on live WT and GOF mice at P80. At P80, total cortical volume in GOF mice is 15% larger than in WT mice, while the volumes of the thalamus and hippocampus remain unchanged (Fig. 2d). Interestingly, the anterior cingulate cortex, which makes up 3% of the total cortical volume, is reduced by 15% in GOF mice, mirroring findings in ASD patients^35, 36^. Nissl staining of brains at P10 and P50 reveals that cortical thickness is similar between WT and GOF mice (Fig. 2e). These results suggest that the increased cortical volume observed in MRI is primarily due to tangential expansion of cortical surface area rather than radial expansion of cortical thickness.

**Figure 2.**
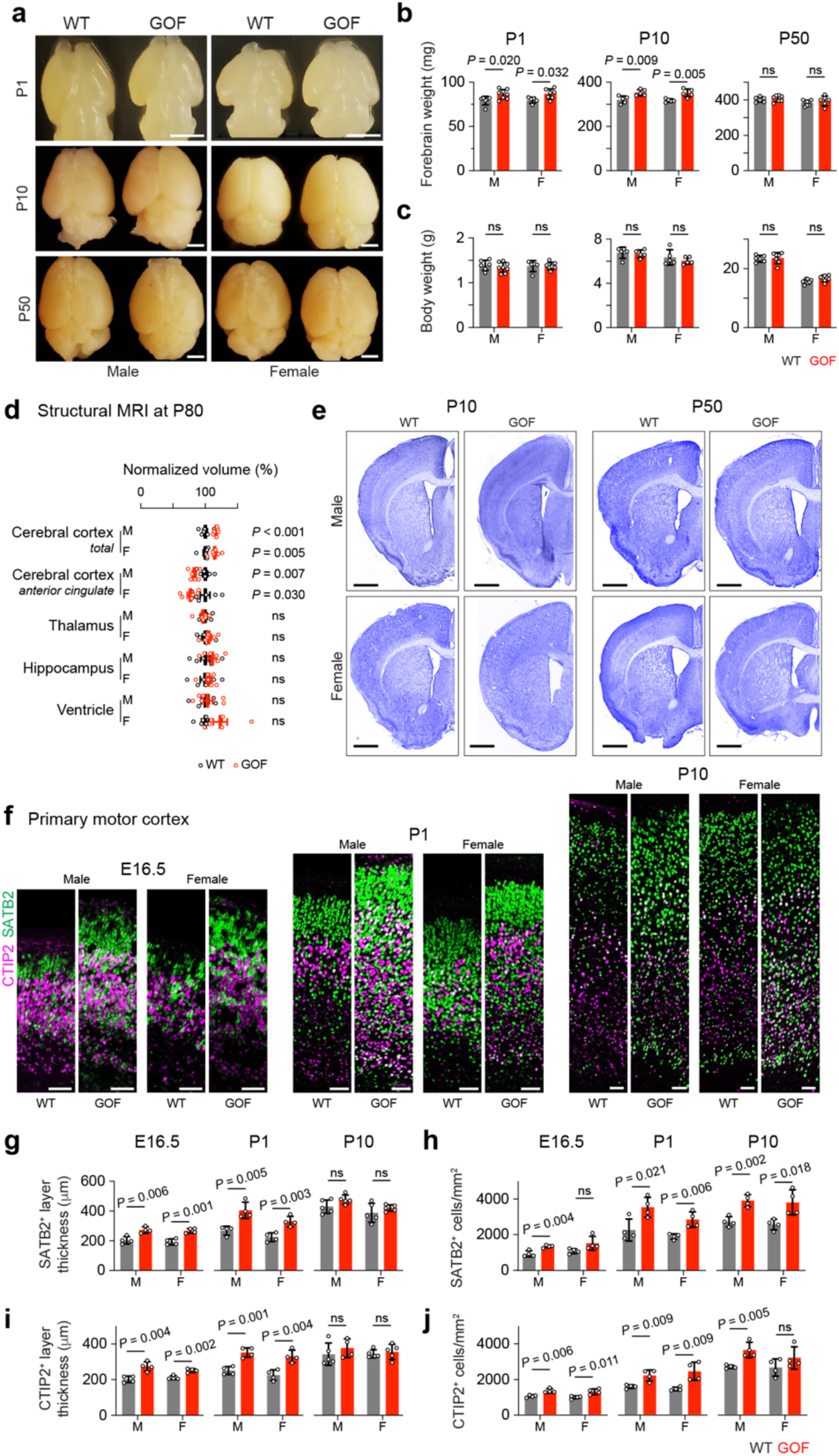
*Aspm* GOF mice are macrocephalic with higher cortical neuron density. **(a-c)** Postnatal *Aspm* GOF mice exhibit macrocephaly as early as P1. Brain weight is significantly increased in both sexes at P1 and P10, while body weight remains unaffected. By P50, brain weight measurement is insufficient to detect cortical volume expansion. N = 5-8 samples/genotype/sex. Student’s *t*-test. Scale bars, 1 mm. **(d)** Structural MRI reveals a significant and sustained increase in total cerebral cortical volume, along with a decrease in the volume of the anterior cingulate cortex at P80. N = 6 mice/genotype/sex. Student’s *t*-test. **(e)** Nissl staining shows similar cortical thickness between WT and GOF mice at P10 and P50. N = 3 mice/genotype/sex. Scale bars, 1 mm. **(f-j)** Immunostaining for the upper-layer marker SATB2 and the deep-layer marker CTIP2 in the primary motor cortex shows thicker cortical layers in GOF at E16.5 and P1, but not at P10. In contrast, increased neuronal density is observed at all three ages examined. N = 4 mice/genotype/sex. Student’s *t*-test. Error bars, mean ± s.e.m. Scale bars, 50 μm

We further examined the cortical structure by immunostaining for the cortical upper-layer neuronal marker SATB2 (layers 2-5) and the deep-layer neuronal marker CTIP2/BCL11B (layers 5, 6, and subplate) (Figs. 2f-2j). At E16.5 and P1, the SATB2^+^ layers and CTIP2^+^ layers are significantly thicker in GOF mice compared to WT mice; however, this difference disappears by P10. Notably, we observed that the density of SATB2^+^ neurons is significantly higher in GOF mice than in WT mice across all three stages examined. The density of CTIP2^+^ neurons is also higher in GOF mice, though this increase was less pronounced in P10 GOF female mice (Fig. 2j). Taken together, these data demonstrate that the *ASPM* GOF mutation induces macrocephaly with increased cortical volume and neuronal density, while maintaining comparable cortical thickness in mice.

### *Aspm* GOF mice show excessive embryonic neurogenesis with increased oRG-like progenitors

*ASPM* is a key regulator of cortical progenitor cell proliferation, with LOF mutations leading to progenitor pool depletion and microcephaly. To determine whether the GOF mutation has the opposite effect—enhancing progenitor proliferation and expanding the progenitor pool—we first analyzed the E14.5 mouse cortex by immunostaining the proliferation marker phospho-Histone H3 (pHH3). GOF mice exhibited a significant increase in pHH3^+^ progenitors compared to WT controls (Figs. 3a, b), with this effect being more pronounced in males. Notably, pHH3^+^ progenitors in the subventricular zone (highlighted by arrowheads), indicative of outer radial glia oRG-like cells^37^, are dramatically elevated in male GOF mice.

**Figure 3.**
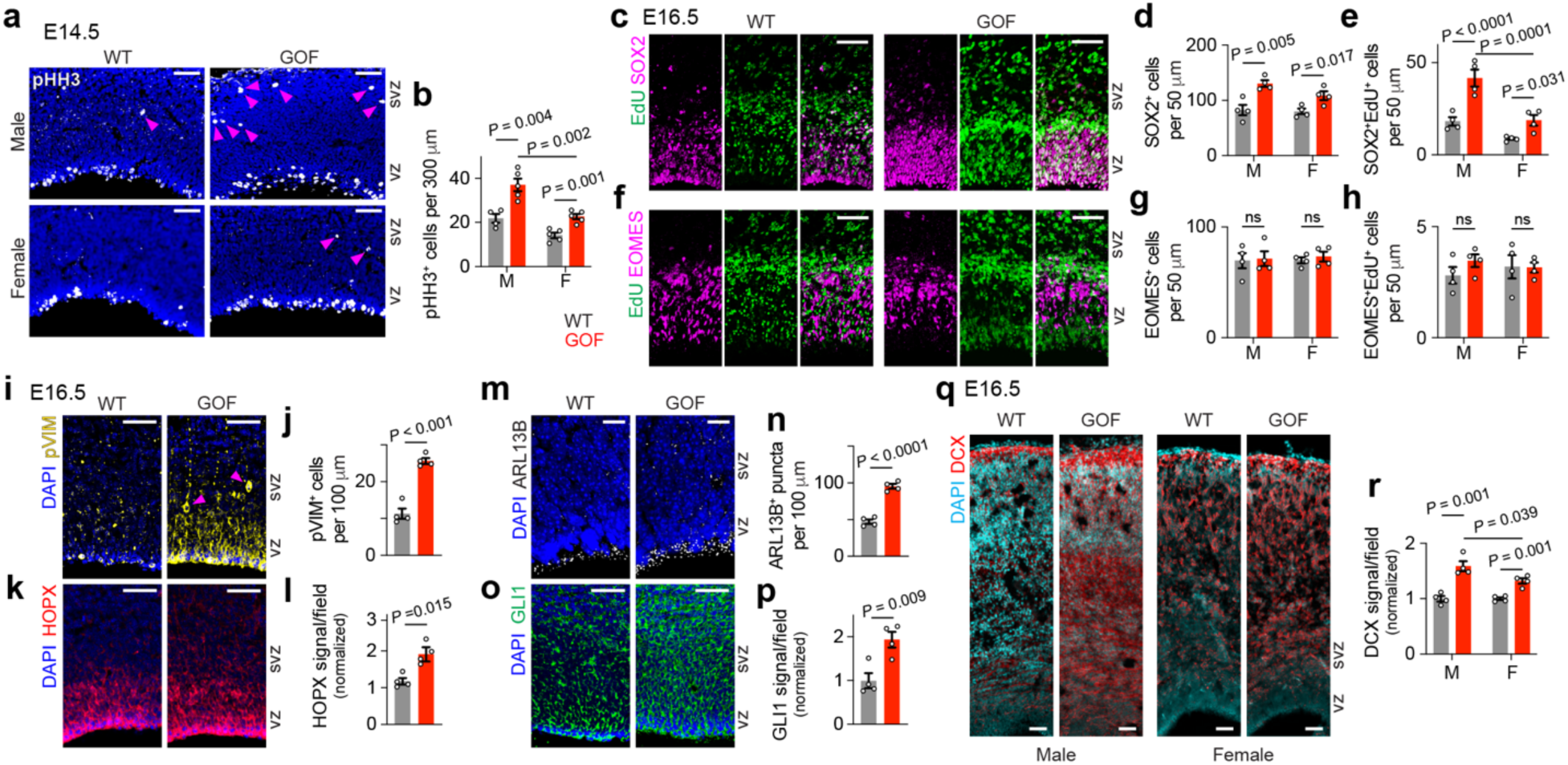
*Aspm* GOF mice show excessive embryonic neurogenesis with increased oRG-like progenitors. **(a, b)** The number of pHH3^+^ progenitor cells in the ventricular zone (vz) and subventricular zone (svz) is increased in GOF mice at E14.5, with a more pronounced increase in males. Arrowheads indicate oRG-like cells in the subventricular zone, which are increased in GOF mice. **(c-e)** At E16.5, the number of SOX2^+^ vRG is increased in GOF mice, with a more pronounced increase in the number of SOX2^+^EdU^+^ proliferating vRG in male GOF mice. **(f-h)** The number of EOMES^+^ and EOMES^+^EdU^+^ IPs is unaffected in GOF mice. **(i-k)** RG cells positive for the proliferation marker pVIM and the oRG/vRG marker HOPX immunostaining signal are increased in E16.5 male GOF mice. Arrowheads indicate oRG-like cells in the subventricular zone. **(m-p)** The number of ARL13B^+^ cilia and Hedgehog downstream GLI1 signal are increased in E16.5 male GOF mice. Notably, ARL13B^+^ puncta are also increased in the subventricular zone of GOF mice. **(q,r)** The immature, migrating neuronal marker DCX is increased in both male and female GOF mice, with a more pronounced increase in males. N = 4 mice/genotype/sex for analyses including both sexes, or 4 mice/genotype for male-only analyses. Student’s *t*-test. Error bars, mean ± s.e.m. Scale bars, 50 μm in all panels, except in panel **(m)**, where scale bars represent 10 μm.

To further characterize the impact of the *Aspm* GOF mutation on cortical progenitor populations, we analyzed the two major progenitor types: SOX2^+^ ventricular radial glia (vRG) and EOMES/TBR2+ intermediate progenitors (IPs). At E16.5, GOF mice exhibit a significant increase in SOX2^+^ vRG compared to WT mice (Figs. 3c-e). To specifically quantify actively proliferating vRG, we administered 5-ethynyl-2’-deoxyuridine (EdU) at E15.5 and harvested embryos 24 hours later. Immunostaining revealed a pronounced increase in SOX2^+^EdU^+^ proliferating vRG in GOF mice at E16.5, with a greater effect observed in males. In contrast, the number of EOMES^+^ IPs remained unchanged in Aspm GOF mice (Figs. 3f-h).

The robust expansion of radial glia in male GOF mice was further validated through immunostaining for phospho-Vimentin (pVIM), a marker of mitotic radial glia, and HOPX, a key marker for both oRG and vRG (Figs. 3i–l). In particular, pVIM^+^ oRG-like progenitors are significantly more abundant in the subventricular zone of *Aspm* GOF mice (Fig. 3i).

Given that ASPM localizes to the centrosome, which serves as the basal body for primary cilia during interphase, and that *Aspm* LOF mice exhibit a reduced number of cilia^15^, we investigated ciliogenesis in *Aspm* GOF mice. Immunostaining for the ciliary marker ARL13B revealed that the number of cilia is doubled in both the ventricular zone and subventricular zone of GOF mice (Figs. 3m, n). Since primary cilia mediate Hedgehog signaling, which controls oRG abundance^38^, we examined GLI1, a key downstream effector of Hedgehog signaling. Consistent with increased ciliogenesis, *Aspm* GOF mice exhibited a significant upregulation of GLI1 expression (Figs. 3o, p). Finally, the expansion of the progenitor pool in *Aspm* GOF mice leads to a significant increase in DCX^+^ immature neurons in both sexes (Figs. 3q, r).

These results demonstrate that *Aspm* GOF mice exhibit excessive embryonic neurogenesis, characterized by an expansion of vRG and oRG-like progenitors, while IPs are unaffected. Importantly, these effects are more pronounced in male mice.

### scRNA-seq reveals increased E-I ratio and differential gene expression in *Aspm* GOF mice

To gain a more comprehensive understanding of the molecular and cellular changes caused by the *ASPM* GOF mutation, we performed single-cell RNA sequencing (scRNA-seq) during embryonic neurogenesis in mice. At E15.5, we collected brain tissues from 3 WT and 3 GOF mice, dissociated them into single-cell suspensions, and processed 30,000-37,000 cells per sample using 10×Genomics Chromium GEM-X Single Cell 3’ v4 platform. This approach yielded high-quality scRNA-seq data, with an average of 34,000 reads, 6,900 unique molecular identifiers (UMIs), and 2,600 detected genes per cell (Supplementary Table 2). For downstream analysis, we selected cells with ≥5,000 UMIs and ≥5% mitochondrial content, resulting in 16,000-18,000 high-quality cells with 2,500-6,000 detected genes/cell.

To classify cell populations, we performed cluster analysis^39^ after Harmony integration of WT and GOF datasets. This analysis identified 22 distinct clusters, each defined by a unique combination of marker gene expression (Fig. 4, Supplementary Fig. 1). Radial glia segregate into 3 clusters based on their differentiation status: RG1 (the most undifferentiated), RG2 (neurogenic), and RG3 (gliogenic). Intermediate progenitors form 2 clusters: IP1 (proliferative) and IP2 (differentiative). Additionally, excitatory neurons in deep layers (ExN-DL) and upper layers (ExN-UL), as well as inhibitory neurons from the medial ganglionic eminence (InN-MGE) and lateral ganglionic eminence (InN-LGE), each form 2 distinct clusters based on their differentiation status. In contrast, inhibitory neuronal progenitors (InNPC) segregate into 3 clusters based on the cell cycle phase (InNPC-G1/S, InNPC-S, and InNPC-M). Meanwhile, hypothalamic (HY), cerebellar (CB), Cajal-Retzius (CR), microglial (MG), and endothelial cells (EC) each form a single cluster. Importantly, *Aspm* is highly expressed in the undifferentiated RG1, proliferative IP2, and InNPC in S and M phases, aligning with previously reported *in situ* hybridization and immunohistochemistry data^12, 15^, validating our cluster analysis (Figs. 4a, b).

**Figure 4.**
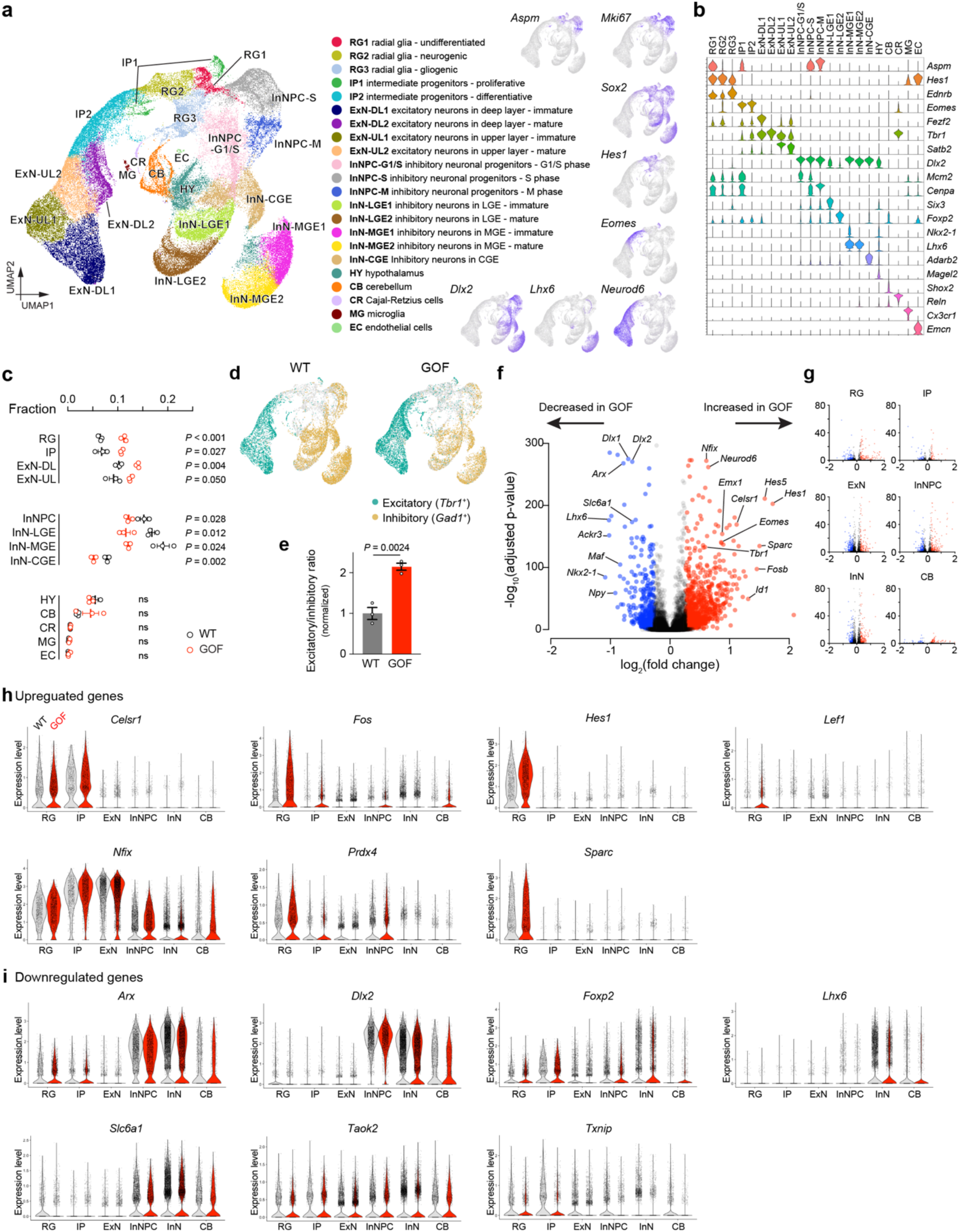
scRNA-seq reveals increased E-I ratio and DEGs in *Aspm* GOF mice. **(a-b)** scRNA-seq analysis of E15.5 WT and *Aspm* GOF mouse brains identifies 22 distinct cell clusters representing vairous lineages, differentiation states, and cell cycle phases. *Aspm* is highly expressed in RG1 (the most undifferentiated radial glia), IP1 (proliferative intermediate progenitors), InNPC-S, and InNPC-M (inhibitory neuronal progenitor cells in S and M phases), which corresponds to the UMAP expression patterns of the proliferation marker *Mki67*. **(c-e)** The proportion of excitatory lineage cells (RG1/2/3, IP1/2, ExN-DL1/2, ExN-UL1/2) is increased, while the proportion of inhibitory lineage cells (InNPC-G1/S/M, InN-LGE1/2, InN-MGE1/2, InN-CGE) is decreased in GOF mouse brains, indicating an increased E-I ratio. The ratio of *Tbr1*^+^ excitatory neurons to *Gad1*^+^ inhibitory neurons is also increased in GOF mice. N = 3 samples/genotype. Student’s t-test. Error bars represent mean ± s.e.m. **(f, h)** DEG analysis of all cell types combined reveals >800 upregulated genes (red) and >500 downregulated genes (blue), mostly reflecting an increased E-I ratio, with some genes, such as *Hes1* in RG, also showing increased expression levels. DEGs also include ASD risk genes such as *Slc6a1* and *Foxp2*. **(g)** DEGs are identified across cell types, irrespective of whether *Aspm* is expressed in the respective cell type.

Surprisingly, we observed a marked increase in the proportion of cells in the excitatory neuronal lineage (RG, IP, ExN-DL, and ExN-UL) and a corresponding decrease in the inhibitory lineage (InNPC, InN-LGE/MGE/CGE) in E15.5 GOF mice, resulting in an increased E-I ratio (Fig. 4c). To further quantify this imbalance, we counted *Tbr1*^+^ excitatory neurons and *Gad1*^+^ inhibitory neurons and observed a 2-fold increase in the E-I ratio in GOF mice (Figs. 4d, e). These results are novel ecause *ASPM* is expressed in progenitors of both excitatory and inhibitory lineages, and its role in regulating the E-I ratio has not been previously reported.

Next, we examined the extent to which the *ASPM* GOF mutation alters gene expression. It turned out that the WT samples consisted of 3 male embryos, whereas the GOF samples included 1 male and 2 female embryos. To isolate the effect of genotype and eliminate potential sex-related variations, we restricted our differentially expressed gene (DEG) analysis to a comparison between one male WT and one male GOF sample. Additionally, we focused on genes expressed in 25% of cells within all or specific clusters, applying a threshold of ≥1.2-fold change with an adjusted p-value <0.05. Under these criteria, we observe 899 upregulated and 511 downregulated DEGs when all clusters are combined (Fig. 4f). Cell type-specific analysis of DEGs (Fig. 4g) revealed changes not only in radial glia and inhibitory neuronal progenitors, where *Aspm* is highly expressed, but also in excitatory and inhibitory neurons, which lack *Aspm* expression. This suggests that the *Aspm* GOF mutation broadly affects gene expression across diverse cell types. Upregulated DEGs are associated with the excitatory neuronal lineage (*Emx1*, *Eomes*, *Neurod6*), progenitor differentiation (*Nfix*), planar cell polarity (*Celsr1*), extracellular matrix remodeling (*Sparc*), immediate early genes (*Fos*, *Fosb*), oxidative stress regulation (*Prdx4*), Bone Morphogenetic Protein (BMP) signaling (*Id1*), Wnt/β-catenin signaling (*Lef1*), and Notch signaling (*Hes1*, *Hes 5*). In contrast, downregulated DEGs are associated with the inhibitory neuronal lineage (*Ackr3*, *Arx*, *Dlx1*, *Dlx2*, *Lhx6*, *Nkx2-1*, *Npy*, *Txnip*), GABAergic signaling (*Slc6a1*), oxidative stress regulation (*Txnip*) and ASD (*Dlx1*, *Dlx2*, *Foxp2*, *Slc6a1*, *Taok2*). Further analysis of individual gene violin plots indicated that the DEGs primarily reflect shifts in the proportions of excitatory versus inhibitory lineage cells, although some genes, especially *Hes1*, also showed sharply increased expression within individual radial glia (Figs. 4h, i).

Our scRNA-seq analysis provides two mechanistic insights. First, the *Aspm* GOF mutation significantly shifts the E-I ratio by disproportionately expanding the excitatory progenitor pool. *Hes1* upregulation in RG may play a crucial role, as a recent study^40^ showed that *Hes1* transgenic mice exhibit vRG and oRG expansion, but not IP expansion, similar to *Aspm* GOF mice. Second, although *Aspm* expression is restricted to progenitors, the GOF mutation drives widespread differential gene expression across major cell types, including the downregulation of ASD-associated genes such as *Foxp2* and *Slc6a1*.

### *Aspm* GOF mice display abnormal neuronal morphology and increased brain connectivity

Postmortem analyses of cortical neurons in individuals with ASD reveal abnormal neuronal arborization and altered synaptic spine density^41^. As *Aspm* GOF mice exhibit increased cortical neuron density (Fig. 2g) and an elevated E-I ratio (Fig. 4), we hypothesized that their neuronal morphology may also be disrupted. Thus, we performed Golgi staining on P50 WT and *Aspm* GOF mice. In the primary motor cortex, Golgi-stained neurons appear more densely packed in GOF mice compared to WT. Analysis of individual pyramidal neurons revealed shorter dendritic length, fewer dendrites, and reduced dendritic branching. In contrast, dendritic spine density is significantly increased in GOF mice (Figs. 5a-g). These results suggest that pyramidal neurons derived from an overly proliferative RG pool in *Aspm* GOF mice show morphological abnormalities, potentially reflecting a compensatory mechanism to enhance synaptic connectivity or a state of dendritic immaturity.

**Figure 5.**
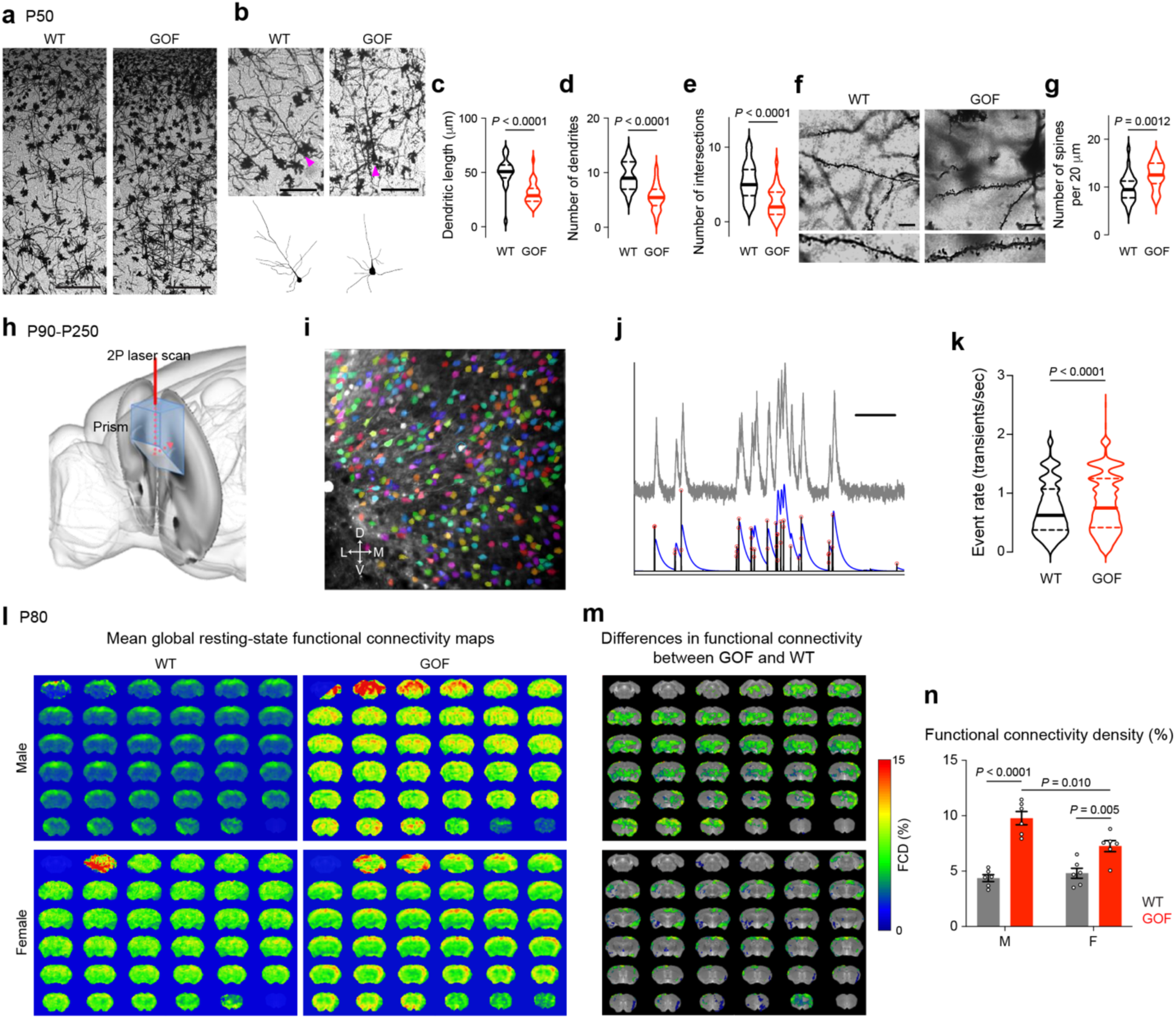
Abnormal neuronal morphology and hyperconnectivity in *Aspm* GOF mouse brains. **(a-g)** Golgi staining of the primary motor cortex at P50 reveals that male *Aspm* GOF mice have a higher density of pyramidal neurons and astrocytes compared to WT mice. GOF neurons exhibit shorter and fewer dendrites with fewer intersections. In contrast, dendritic spine density is increased in GOF neurons. N = 25-30 samples/genotype. Student’s t-test. Scale bars, 100 μm (a) and 10 μm (b, f). **(h)** Schematic of the imaging preparation in Ca^2+^ imaging. Mice (N = 4 WT, 5 GOF) were injected in the prefrontal cortex with AAV encoding GCaMP6f under the Synapsin promoter, and a chronic imaging prism was implanted. **(i)** Mean imaging frame from a single session (Mouse 212, N = 367 identified cells, Suite2P). **(j)** Example fluorescence trace (gray), denoised Ca^2+^ trace (blue), deconvolved trace (black), and thresholded events (red circles). Scale bar = 60 seconds. **(k)** Event rates for GOF (N = 1468 ROIs in 4 mice, median = 0.75 Hz) and WT (N = 3369 ROIs in 5 mice, median = 0.63 Hz, rank sum p = 5 x 10^-11^). **(l)** Resting-state fMRI-based average functional connectivity density (FCD) maps across WT and GOF mice at P80. N = 6/genotype/sex. **(m)** FCD differences between GOF and WT and maps of significantly changed FCDs (p < 0.05) FDR corrected, local cluster size k > 25 voxels, determined by two-tailed *t*-test corrected for multiple comparisons. **(n)** Local functional connectivity in males and females, N = 6 mice/genotype/sex. Student’s *t*-test comparisons for different conditions with Benjamini-Hochberg correction for multiple comparisons with a false discovery rate (FDR) < 0.05 and local cluster size of k > 25 voxels. Error bars, mean ± s.e.m.

Having identified an increased E-I ratio in GOF mice as measured by relative counts of excitatory and inhibitory neurons, we next examined activity in cortical neurons in these mice, hypothesizing that GOF mice, with greater E-I ratio, would exhibit elevated activity in cortical neurons. We focused our recording on the prefrontal cortex (PFC) for several reasons. First, PFC is critically important for mediating responses to social cues in both humans^42^ and mice^43^, and GOF mice show deficits in social behavior (Fig. 6). Second, structural MRI scans of GOF mice show greatest reduction in cortical volume in the frontal anterior cingulate cortex (Fig. 2d), possibly reflecting a corresponding dysregulation of frontal activity. We therefore performed 2-photon Ca^2+^ imaging of pan-neuronal prefrontal activity in awake WT and GOF mice (4 male WT mice, 5 male GOF mice) between P90 and P250. In line with scRNA-seq data, GOF mice showed elevated activity compared with WT (N = 1468 ROIs in 4 WT mice, median = 0.75 Hz; N = 3369 ROIs in 5 GOF mice, median = 0.63 Hz, rank sum p = 5 x 10^-11^) (Fig. 5h-k).

**Figure 6.**
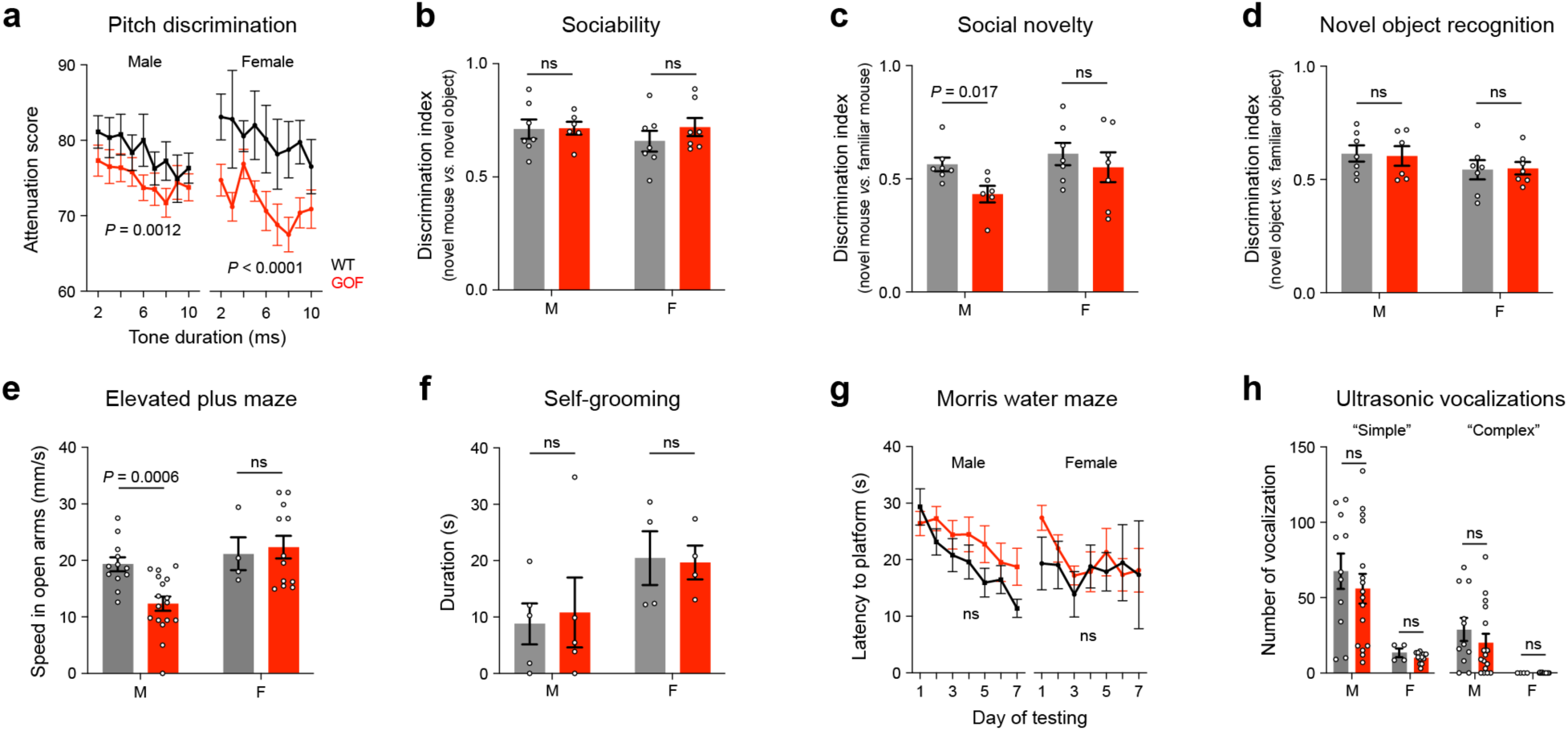
Social-cognitive deficits in *Aspm* GOF mice. **(a)** *Aspm* GOF mice exhibit enhanced pitch discrimination sensitivity compared to WT mice. **(b)** In the three-chamber assay, GOF mice display normal sociability, preferring a mouse over an object; **(c)** however, male GOF mice show deficits in social novelty, failing to prefer a novel mouse over a familiar one. **(d)** GOF mice demonstrate normal novel object recognition, favoring a novel object over a familiar one. **(e)** In the elevated plus maze, male GOF mice exhibit reduced movement speed in the open arms compared to male WT and female mice. **(f-h)** GOF mice show no impairments in repetitive self-grooming behavior, spatial learning and memory (Morris water maze), or vocal communication (ultrasonic vocalizations). For pitch discrimination, elevated plus maze, Morris water maze, and ultrasonic vocalization experiments, 12 WT male, 17 GOF male, 4 WT female, 12 GOF female mice were used in two cohorts between P80 and P200. For sociality, social novelty, novel object recognition, and self-grooming, 5-7 WT male, 5-6 GOF male, 4-7 WT female, 4-7 GOF female mice were used at P35. Statistical significance was calculated using two-way or three-way ANOVA. Error bars, mean ± s.d. for pitch discrimination and Morris water maze, and mean ± s.e.m. for other assays.

Building on the findings of increased dendritic spine density and elevated Ca^2+^ activity in the PFC of *Aspm* GOF mice, we next examined local functional connectivity across the entire brain. To achieve this, we utilized resting-state functional MRI (rsfMRI), a technique that generates whole-brain voxel maps corrected for multiple comparisons, allowing for the unbiased identification of brain regions with abnormal local functional connectivity^44^ (Figs. 5l-n). In P80 *Aspm* GOF mice, we observed a marked increase in global resting-state functional connectivity across both cortical and subcortical regions. The increase is significant in both males and females but is more pronounced in male mice.

Together, our integrative analysis using Golgi staining, Ca^2+^ imaging, and rsfMRI provides compelling evidence that *Aspm* GOF mice exhibit hyperconnectivity in the PFC, multiple cortical regions, and subcortical areas across various postnatal stages. These results align with observations in human ASD patients^45^, further supporting the relevance of *Aspm* GOF mice as a model for studying ASD-associated neural circuit abnormalities.

### *Aspm* GOF mice show social-cognitive deficits

To explore the functional consequences of macrocephaly in ASD at the organismal level, we conducted behavioral assays in *Aspm* GOF mice and age-and sex-matched littermates between P35 and P200. This approach serves as a critical test to determine whether macrocephaly alone can drive behavioral symptoms associated with ASD. As sensory hypersensitivity, particularly to auditory stimuli^46^, affects 50–70% of individuals with ASD, we first evaluated auditory function in *Aspm* GOF mice. We utilized the variable duration embedded tone test^47^, a paradigm previously applied to detect auditory hypersensitivity in the *Cntnap2* knockout ASD mouse model. This test assesses an animal’s ability to detect a frequency shift from a 75-dB, 10,500-Hz continuous background tone to a 75-dB, 5,600-Hz pure-tone stimulus, with cue durations ranging from 0 to 10 ms. Both male and female Aspm GOF mice display significantly lower attenuation scores compared to WT mice, indicating enhanced pitch discrimination (Fig. 6a).

Next, we evaluated sociability and social novelty in *Aspm* GOF mice using the three-chamber assay. *Aspm* GOF mice exhibit normal sociability, similar to WT mice, by spending more time with a novel mouse than with a novel object (Fig. 6b). However, male *Aspm* GOF mice showed a significant deficit in social novelty preference (Fig. 6c). When given the choice between a novel and a familiar mouse, male GOF mice spend significantly more time with the familiar mouse, a pattern distinct from male WT mice. Notably, this deficit in social novelty preference is not observed in female GOF mice. Such impairments in social novelty preference have also been reported in other ASD mouse models lacking *Chd8*^48^, 16p11.2^49^, and *Shank3^50^*. In contrast, *Aspm* GOF mice display normal novel object recognition, preferring a novel object over a familiar one (Fig. 6d). This indicates that the preference for novelty is specifically impaired in social interactions with conspecifics, rather than with objects, highlighting the social deficits in *Aspm* GOF mice.

Anxiety is a common comorbidity observed in 50% of patients with ASD^51^. To assess anxiety, we performed the elevated plus maze assay, where a mouse explores both open and closed arms. We found that male *Aspm* GOF mice, but not female GOF mice, significantly reduce their exploration time and speed in the open arms (Fig. 6e). This behavior likely reflects increased anxiety rather than motor deficits, as male GOF mice exhibit similar activity levels and movement speeds to WT mice in the open field test (data not shown). Furthermore, we observed no differences in self-grooming (a measure of repetitive behavior), Morris water maze (spatial learning and memory), or ultrasonic vocalizations (vocal communication), further emphasizing the specificity of the behavioral deficits in *Aspm* GOF mice.

In summary, the behavioral assays revealed that *Aspm* GOF mice exhibit auditory hypersensitivity (without a sex difference), social novelty deficits, and anxiety, with the latter two being male-specific. These findings suggest that Aspm GOF mice display a subset of ASD-related behavioral symptoms that may be attributed to macrocephaly, with sex-specific differences in certain behaviors.

## DISCUSSION

In this study, we aimed to determine the role of macrocephaly in the pathogenesis of ASD. Our approach involved using the GOF mutation in *ASPM* (A1877T in humans and A1845T in mice) to induce macrocephaly by expanding the cortical progenitor pool without directly affecting the terminally differentiated neurons. ASPM is known to regulate the proliferation of cortical progenitor cells during embryonic development, and this GOF mutation leads to increased ASPM protein levels. *Aspm* GOF KI mice, which carry the orthologous mutation, exhibit macrocephaly characterized by an expansion in cortical volume and increased neuronal density, though cortical thickness remains unchanged. The *Aspm* GOF mutation specifically increases the progenitor pool of vRG, but not IPs. Unexpectedly, the mutation also results in a significant increase in the number of oRG-like progenitors in the subventricular zone. scRNA-seq analysis at E15.5 reveals that, although *Aspm* is expressed in both excitatory and inhibitory neuronal progenitors, the GOF mutation disproportionately increases the excitatory progenitor pool. This shift results in a marked increase in the E-I ratio, likely through the upregulation of *Hes1*^40^ and other DEGs associated with neurogenesis and ASD. Further analyses, including Golgi staining, calcium imaging, and rsfMRI, reveal defects in neuronal morphology and brain-wide hyperconnectivity, which exhibit male-specific effects. Behaviorally, *Aspm* GOF mice demonstrate auditory hypersensitivity, deficits in social novelty preference, and moderate anxiety. These findings strongly suggest that macrocephaly alone can drive a subset of ASD-like symptoms. More importantly, our results emphasize that macrocephaly in the context of ASD is not a proportional expansion of excitatory and inhibitory neurons. Instead, it involves a skewing of the E-I ratio, even though the causative gene is equally expressed in both excitatory and inhibitory progenitors, with no previous studies indicating its role in modulating the E-I ratio.

We propose that the method used to identify the A1877T GOF mutation—the discovery of genetic variants associated with both ASD and cancer through publicly available databases—can be applied to identify GOF mutations in other neurodevelopmental genes. This approach may be particularly valuable for genes in which LOF mutations are linked to microcephaly, such as *CTNNB1*. Because macrocephaly is observed in approximately 20% of ASD patients, it is likely that many genes and signaling pathways are shared between ASD and cancer, providing a basis for the identification of additional relevant genetic variants.

Although the A1877T GOF mutation has significant effects on the cortical progenitor pool, DEGs, brain connectivity, and behavior, the increase in endogenous ASPM protein levels is only 1.5-fold, as measured by Western blot analysis. This modest increase was unexpected but is consistent with recent findings indicating that the impact of GOF mutations on protein structure is considerably milder than that of LOF mutations^52^.

Our research raises two key questions. First, given that macrocephaly caused by the *ASPM* GOF mutation is associated with an increased E-I ratio, is microcephaly caused by the *ASPM* LOF mutation associated with a decreased E-I ratio? In both macrocephalic ASD and microcephaly patients, intellectual disability (ID) is a significant comorbidity. Is the nature of ID different between macrocephaly and microcephaly, particularly in terms of the E-I ratio, or do these conditions share paradoxical similarities? Second, although macrocephaly is observed in both male and female GOF mice, brain hyperconnectivity, social novelty deficits, and anxiety are significantly more pronounced in males. What mechanisms underlie the differential functional consequences of macrocephaly? Future studies are needed to address these questions.

### Limitations

It is unknown whether the seven ASD patients carrying the A1877T heterozygous mutation exhibit macrocephaly. Although genetic studies have identified at least 23 individuals homozygous for this mutation^53^, no clinical data are available. Detailed information on specific genetic variants is often lacking unless they significantly impact the carrier’s quality of life and are documented by physicians. In this context, human organoid studies could provide valuable additional insights.

Our macrocephalic *Aspm* GOF mouse model will provide novel mechanistic insights into ASD and neurodevelopmental disorders in general, especially when studied in parallel with microcephalic *Aspm* LOF mice.

## METHODS

All animal experiments were carried out under protocols approved by the IACUCs of the University of Connecticut School of Medicine (Farmington, CT), the University of Connecticut (Storrs, CT), and Yale University (New Haven, CT).

### Generation and maintenance of *Aspm* GOF mice

To generate *Aspm* GOF KI mice carrying the A1845T (c.5533G>A) mutation, over 100 C57BL/6 zygotes were injected with eSpCas9 protein (MilliporeSigma ESPCAS9PRO) at 100 ng/μl, a crRNA:tracrRNA duplex (TCAATTTCTAAAGACTAGAGAGG) at 40 ng/μl, and a single-stranded DNA oligonucleotide for homology-directed repair (A*C*GTAGACTGGATCTTCACTGCAGCCTCATGTTGCCTCCTGAGCTGTTGCCGAACCTGC CAGCCACGGTAGGCAGACTGGAGGCAAACCACAG<u>T</u>CTCTCTAGTCTTTAGAAATTGAATTCT CATGTCATAAGCAACCTTCTGTGCCCTGTACCATCTCTGAATCTTCACAGCAGACTGAAGCA TAGTCTGGTATTTCA*T*C, *phosphorothioate modification) at 50 ng/μl with phosphorothioate modifications. Microinjections were performed at the UConn Center for Mouse Genome Modification, and the injected zygotes were implanted into the oviducts of pseudopregnant females and carried to term. Two male *Aspm* GOF KI founders heterozygous for the targeted mutation were identified by Sanger sequencing of the modified locus and were bred with WT C57BL/6 females to confirm germline transmission. Genotyping was performed using PCR with 4 primers that distinguish the WT (227-bp) and GOF (194-bp) alleles (GCAGGCAGCTTACAGAGGC, CCTCACATGCTGCTGGATGAC, GAATTCAATTTCTAAAGACTAGAGCGA, CAGACTGGAGGCAAACCACCGC). Both heterozygous and homozygous *Aspm* GOF KI mice were viable, fertile, and maintained through crossbreeding with WT C57BL/6 mice. Sex determination of postnatal mice was performed by visual examination, while embryonic sex was identified using PCR primers targeting an 84-bp deletion in the X-linked *Rbm31x* gene relative to its Y-linked gametolog *Rbm31y*, producing a single 269-bp amplicon in females (XX) and both 269-bp (X) and 353-bp (Y) amplicons in males (XY), as previously described^55^.

### Alignment of vertebrate ASPM proteins

Based on RefSeq protein numbers at NCBI Orthologs, we aligned protein sequences from 8 species using EMBL multiple sequence comparison by log-expectation (MUSCLE): human (NP_060606.3), macaque (XP_045250348.2), marmoset (XP_002760652.3), mouse (NP_033921.3), ferret (XP_044942926.1), pig (XP_020920137.1), chicken (XP_025009015.2), and zebrafish (XP_005161794.1).

### Immunostaining and Western blot analysis

We performed immunostaining and Western blot analysis of mouse brain samples as described previously^15^ using the following antibodies: ASPM (rabbit, 1:100, gift from W. Huttner), ARL13B (rabbit, 1:200, Proteintech). beta-actin (rabbit, 1:2000, Proteintech), Centrin (mouse, 1:1000, Millipore), CTIP2/μ11B (rat, 1:100, Millipore), DCX (rabbit, 1:100, ThermoFisher; rabbit, 1:500, Abcam), EOMES (rabbit, 1:100, Genetex), GFP (chicken, 1:100, Abcam), GLI1 (rabbit, 1:100, Novus), HOPX (mouse, 1:100, Santa Cruz), pHH3 (mouse, 1:200, Proteintech), pVIM (mouse, 1:200, MBL), SATB2 (rabbit, 1:100, Bethyl), SOX2 (mouse, 1:50, Santa Cruz). Images were acquired by white field LSM-700 confocal microscope (Carl Zeiss, Germany) and analyzed by ImageJ software (NIH, USA).

### Golgi staining

Golgi staining was performed as per the manufacturer’s instructions (FD Neurotechnology, MD, USA) with minor modifications. Under isoflurane anesthesia, mice were transcardially perfused with ice cold PBS followed by 4% PFA and incubated in 4% PFA for overnight at 4°C. The next day, brains were incubated in an impregnation solution for 72 h in the dark at 37°C. Impregnation solution was prepared by mixing equal volumes of solution A and solution B provided in the kit and replaced at every 24 h. After 72 h, brains were incubated in solution C (provided in the kit) for a total of 48 h at 4°C in the dark, and solution was replaced with fresh solution after 24 h. Brains were cut into 80 µm thickness coronal sections using a cryostat (Leica, Germany) and mounted on gelatin-coated Superfrost Plus slides (Fisher Scientific). Sections were placed in freshly prepared staining solution provided in the kit for 10 min at room temperature. Sections were washed with water and dehydrated in sequential rinses of 50%, 75%, 95% and 100% ethanol 4 min each rinse. Sections were cleared with 3 times rinse in xylene and mounted with Permount mounting media (Fisher Scientific, USA). Sections were imaged on 10x magnification for the analysis of neuron distribution and density in frontal cortex. For the analysis of dendritic length, number of intersections and spine density, sections were imaged at 63x magnification. Images were acquired by white field LSM-700 confocal microscope (Carl Zeiss, Germany) and analyzed by ImageJ software (NIH, USA).

### scRNA-seq analysis

Three WT and three *Aspm* GOF embryonic mouse brains were collected from time-pregnant females, dissociated into single cells using Neural Tissue Dissociation Kit (Miltenyi Biotec), and processed the JAX-UConn Single Cell Genomics Center (Farmington, CT). Each sample yielded 30,000-37,000 cells, which were sequenced on the 10ξGenomics Chromium GEM-X Single Cell 3’ v4 platform, generating an average of 34,000 reads, 6,900 UMIs, and 2,600 detected genes per cell (Supplementary Table 2). For downstream analysis, we selected cells with ≥5,000 UMIs and ≥5% mitochondrial content, resulting in 16,000-18,000 high-quality cells with 2,500-6,000 detected genes/cell. Data processing, cell clustering, and data analysis were performed as previously described^39^. For DEG analysis, we focused on genes expressed in ≥5% of cells. Genes with an average fold change ≥1.2 and p-value ≤0.05 were considered significantly altered.

### Calcium imaging

#### Surgery

Animals (*Aspm* GOF or WT littermates 6-10 weeks) were placed inside a flow box and anesthetized with isoflurane gas (2%) until sedated, at which point they were placed in a stereotax and maintained on 0.5% isoflurane for the duration of the surgery. Scalp hair was trimmed away, and a midline incision was made using fine surgical scissors (Fine Science Tools), exposing the skull. The periosteum was bluntly dissected away and bupivacaine (0.05 mL, 5 mg/kg) was topically applied. For prism implantation, a large rectangular craniotomy was made above left PFC, extending from 1.5mm anterior to 3.7mm anterior, and from 2.0mm lateral (left) to 0.2mm lateral (right, across midline).

A 0.5-mm burr (Fine Science Tools) and a high-speed hand dental drill (Osada) were used, taking great care not to compress brain tissue or damage the sagittal venous sinus. Gentle irrigation with phosphate-buffered saline (137 mM NaCl, 27 mM KCl, 10 mM phosphate buffer, VWR) was used to clear debris at regular intervals. The dura beneath the craniotomy was removed using fine forceps (Fine Science Tools).

Chronically implanted microprisms were 1.5mm X 1.5mm X 3mm microprism (M/L,A/P,D/V) from OptoSigma (BK7 borosilicate glass with aluminum hypotenuse and silicon dioxide coating), which was implanted at a depth of 2.3mm ventral to brain surface using a stereotaxic micromanipulator (Kopf), with the prims held in place using vacuum suction via an 18G blunt needle. Minimal reactive gliosis was seen in the coronal imaging field, and maximum calcium-mediated fluorescence was seen 50-150μM past the prism face, and therefore imaging planes were confined to this depth.

All head-fixed animals received custom-machines stainless steel head plates affixed with Metabond dental cement. Head plates featured a circular central aperture centered around the imaging field (9mm I.D.), with right and left arms securing (25mm total width) that accommodated 0-80 socket screws (0.38g in total). Sterile eye lubricant (Puralube, FischerSci) was administered to prevent corneal drying, and a microwavable heating pad (Snugglesafe) was used to maintain body temperature. Metacam (1 mg/kg, i.p.) was administered after surgery as a prophylactic analgesic.

#### Viral transduction

AAV9 of titer exceeding 1012vg/ml (Addgene) was used to package the plasmids. For imaging experiments, AAV9-hSyn-GCaMP6f was targeted to PFC. Injection coordinates for PFC were 1.75mm anterior, 0.35mm left of midline. Two D/V sites received infusions, at 2.0mm and 1.5mm ventral to brain surface. A Nanoject small volume injection pump (Drummond Scientific) was used to infuse the viral suspension through a pulled glass capillary pipette (WPI), which was allowed to sit 5min to allow for tissue settling before infusion. Virus was infused at a rate of 50nL/min, for a total of 250nL per site.

#### 2-photon imaging

Imaging was performed on a Thorlabs Bergamo II imaging system via a 10X apochromatic objective (air immersion, 0.5NA, 7mm WD). All images were acquired using a Chameleon Discovery NX laser tuned to 920nm. Fluorescence was recorded through gallium arsenide phosphide (GaAsP) detectors using Thorimage acquisition software using a green light emission bandpass filter (Semrock). Three-plane imaging was performed with a distance of 25μm between planes, at a rate of 7.5 volumes per second, and with a pixel resolution of 512 x 512 pixels over 660μm X 660μm field of view, for a μM/pixel ratio of 1.28. All calcium imaging experiments occurred in awake mice.

#### Image processing

Videos were processed using Suite2P^56^, including frame registration and ROI detection. To detrend low-frequency shifts in signal intensity, a three-step procedure was used. First, neuropil signal was subtracted from the ROI fluorescence signal. Second, shifts in calcium transient amplitude due to, e.g., bleaching and tissue drift were corrected by identifying signal peaks (Matlab, findpeaks) and fitting a second-order polynomial function to peak amplitudes, then multiplying the fluorescence signal by the inverse of this function. Third, baseline drift was detrended by masking transient times (performing linear interpolation through epochs in which the signal exceeded 3 median absolute deviations), subtracting a Savitzky-Golay-filtered signal from this masked signal, then unmasking the transient epochs by adding the original fluorescence signal from these times to the detrended baseline. Constrained FOOPSI^57^ was then used to denoise the signal and produce a deconvolved calcium event trace. Event rates were derived from this deconvolved trace by detecting peaks (Matlab findpeaks) and computing the inverse of the median inter-event interval across the recording session.

### Resting state fMRI (rsfMRI)

We used functional connectivity density (FCD) mapping to assess the local functional connectivity. FCD is a voxel- and degree-based metric, which identifies the number of correlated voxels to a base voxel without identifying the precise location of the correlated voxels. These metrics are degree based, as they are based on the number of voxels that a given voxel is strongly correlated with, or functionally “connected to.” Global FCD determination is a standard graph theory-based analysis determining brain functional connectivity using resting state fMRI and originally demonstrated in the human brain^58^. Local functional connectivity was determined as described previously^59, 60^. Mice were initially anesthetized with 2-3 % isoflurane, and a PE 50 tubing was placed into the i.p. cavity for dexmedetomidine (sedative) infusion. Thereafter, mice were maintained under complete anesthesia with 0.25% isoflurane and 250 μg/kg/h i.p, of dexmedetomidine. Body temperature was maintained at 36°C–37°C using a heating pad and monitored using an MRI-compatible rectal probe throughout the MRI experiments. Heart rate and respiratory rate were continuously monitored throughout the MRI experiments. Dynamic blood oxygenation dependent (BOLD) data were obtained using a Bruker 9.4T/16 magnet (Bruker BioSpin, MA, USA) using a single-shot gradient echo, echo planar imaging (GE-EPI) sequence with the following parameters: TR of 1000 ms, TE of 12 ms, in-plane resolution of 400 × 400 μm and slice thickness of 1000 μm, for a total of 300 images for each run. BOLD time series across all voxels were detrended using a second order fit and bandpass filtered (0.001 to 0.1 Hz) to exclude slow drift of the signal. Images were then registered to a brain template (200 x 200 x 200 um spatial resolution) followed by spatial Gaussian filtering (FWHM=1.5 mm) as previously described^59, 60^. For statistical analysis, local functional connectivity maps were calculated for each voxel using threshold correlation (T_c_) > 0.6, a signal-to-noise ratio (T_SNR_> 0.5) in individual space and normalized to a z score with the Fisher r to z transformation. Significance was assessed using same-sex Student’s t-test comparisons for different rearing conditions with Benjamini-Hochberg correction for multiple comparisons with a false discovery rate (FDR) < 0.05 and local cluster size of k> 25 voxels.

### Behavioral assay

On day 1 of the battery, 5-week-old male and female mice started to be handled for 3 mins, and it lasted until day 4 to familiarize with the experimenter. The experimenter was blinded to their genotype. For all the behavioral experiments, any apparatus was cleaned with 70% EtOH, distilled water, and dry paper towels in order between animals. All the tests were performed during the light time of the animal facility’s light/dark cycle under dim light. ANY-maze software was used to record and track the location and movement of mice.

#### Three-chamber assay

On day 2 and day 3, the subject mice were habituated to the three-chambered apparatus and two empty wire cups for 10 mins. Age and background-matched stranger mice, which had never met the subject mice, were habituated in a wire cup on day 2 and day 3, being kept separately from the subject mice. On day 4, subject mice were habituated to the apparatus for another 10 mins. Then, a stranger mouse (stranger 1) or a yellow block (object) was introduced to each wire cup, and the movement of the subject mouse was tracked for the next 10 mins. Finally, the yellow block was replaced with another stranger mouse (stranger 2), and the movement of the subject mouse was tracked for the next 10 mins.

#### Open field test

On day 8, the mice were placed in a box (40 x 40 x 40 cm) and their movement was tracked for 10 mins. Center was defined as the square area occupying 50% of the entire space in the middle of the box.

#### Repetitive behaviors

Time spent in repetitive self-grooming and rearing was manually counted from videos of the open field test.

#### Novel object recognition

On day 9, two same objects (old) were placed in two different diagonal corners in the same box which was used for the open field test. The mice were placed in the box for 10 mins. On day 10, one of the objects was changed to a different object (novel) and the mice was placed in the box and their movement was tracked for 5 mins. For objects, 15 ml conical tubes and slide mailers were used and counterbalanced between old and new.

#### Pitch discrimination, elevated plus maze, Morris water maze, and ultrasonic vocalization

We performed these assays as previously described^47, 61^.

### Statistics

All statistical analyses were performed using GraphPad Prism. Error bars represent mean ± SEM unless otherwise specified. Behavioral data were analyzed using the two-tailed Mann-Whitney test, a commonly used non-parametric method for comparing WT and GOF mice.

## ACKNOWLEDGEMENTS

We would like to thank the UConn Health Center for Comparative Medicine, the UConn Animal Care Services, and the Yale Animal Resources Center for professional care of *Aspm* GOF and WT mice. We thank W. Huttner for mouse ASPM antibody, M. Chatterjee for the initial characterization of ASD-associated *ASPM* variants, K. Jorgensen for tracing Golgi-stained neurons, and H. Faraday for mouse husbandry. This work was supported by UConn Institute for the Brain and Cognitive Sciences (B.-I.B), Eagles Autism Foundation (B.-I.B), and National Institute of Child Health and Human Development grant R21HD108696 (B.-I.B).

## AUTHOR CONTRIBUTIONS

B.-I.B. conceived the study and designed and supervised experiments. S.-P.Y. generated *Aspm* GOF KI mice. S.S. performed most of the immunohistochemical and histochemical experiments, and prepared scRNA-seq samples. H.K. performed Western blot analysis, optimized immunostaining condition, and the behavioral assays using P35 mice. A.E. and R.C. carried out behavioral assays using P80-P200 mice and J.S. and R.H.F. supervised. T.S. performed calcium imaging and A.M.L. assisted. B.S., and V.G. performed MRI and F.H. supervised. S.R.P. analyzed scRNA-seq data. J.G. confirmed statistical analysis. S.S. and B.-I.B. wrote the manuscript. All authors provided edits and comments on the manuscript.

---

**Supplementary Table 1.** *ASPM* missense variants associated with ASD and cancer.

| No. | Position | Nucleotide change in ASD | AA change in ASD | Inheritance pattern | Parental transmission | Family type | PubMed ID | Original AA conserved in vertebrates | Cancer cases in COSMIC |
| --- | --- | --- | --- | --- | --- | --- | --- | --- | --- |
| 1 | 490 | c.1468C>T | R490C | Familial | - | - | 28831199 |  |  |
| 2 | 523 | c.1567A>C | S523R | De novo | - | Simplex | 35982159 |  |  |
| 3 | 627 | c.1880G>A | R627H | De novo | - | - | 35982159 |  |  |
| 4 | 663 | C>A | A663S | Familial | Maternal | Simplex | 25961944 | Yes | 1 |
|  | 663 | C>A | A663S | Familial | Paternal | Simplex | 25961944 | Yes |  |
|  | 663 | c.1987G>T | A663S | Familial | Paternal | Simplex | 25961944 | Yes |  |
| 5 | 683 | c.2047C>T | P683S | Familial | Paternal | Simplex | 25961944 | Yes | 1 |
| 6 | 756 | c.2267A>G | Y756C | Familial | Paternal | Simplex | 25961944 | Yes |  |
|  | 756 | c.2267A>G | Y756C | De novo | - | Multiplex (monozygotic twins) | 39038432 |  |  |
| 7 | 759 | c.2275C>T | R759W | Familial | Maternal | Simplex | 25961944 |  |  |
| 8 | 767 | c.2300G>A | R767H | Familial | Paternal | Simplex | 25961944 | Yes | 1 |
| 9 | 780 | c.2339A>C | K780T | Familial | Paternal | Simplex | 25961944 | Yes |  |
| 10 | 785 | c.2353C>T | L785F | De novo | - | Simplex | 37393044 |  |  |
| 11 | 871 | c.2612C>T | P871L | Familial | Maternal | Simplex | 25961944 | Yes | 1 |
| 12 | 874 | c.2621A>G | Y874C | Familial | Maternal | Simplex | 25961944 |  |  |
| 13 | 916 | c.2746G>A | D916N | Familial | Maternal | Simplex | 25961944 | Yes |  |
| 14 | 942 | c.2824C>T | R942C | Familial | Paternal | Simplex | 25961944 | Yes | 4 |
| 15 | 993 | c.2977A>G | K993E | Unknown | - | - | 39342494 |  |  |
| 16 | 1031 | c.3091A>G | I1031V | Familial | Maternal | Simplex | 25961944 | Yes |  |
| 17 | 1132 | c.3395A>T | E1132V | Familial | Maternal | Simplex | 25961944 | Yes |  |
| 18 | 1132 | c.3395A>G | E1132G | Familial | - | - | 28831199 | Yes |  |
| 19 | 1136 | c.3406G>A | V1136M | Familial | Maternal | Simplex | 25961944 | Yes |  |
| 20 | 1166 | c.3497C>T | T1166I | Familial | Maternal | Simplex | 25961944 |  |  |
| 21 | 1208 | c.3624G>T | E1208D | Familial | Maternal | Simplex | 25961944 | Yes |  |
| 22 | 1264 | C>T | R1264H | Familial | Paternal | Simplex | 25961944 |  |  |
|  | 1264 | c.3791G>A | R1264H | Familial | Maternal | Simplex | 25961944 |  |  |
|  | 1264 | c.3791G>A | R1264H | Familial | Paternal | Simplex | 25961944 |  |  |
| 23 | 1277 | c.3829T>C | W1277R | Familial | Maternal | Simplex | 25961944 | Yes |  |
| 24 | 1432 | c.4296A>C | R1432S | Unknown | - | - | 35205252 | Yes |  |
| 25 | 1557 | c.4669T>C | C1557R | De novo | - | Simplex | 25363768 |  |  |
| 26 | 1667 | c.5000G>A | R1667H | Familial | - | - | 28831199 | Yes | 2 |
| 27 | 1877 | C>T | A1877T | Familial | Maternal | Simplex | 25961944 | Yes | 3 |
|  | 1877 | C>T | A1877T | Familial | Paternal | Simplex | 25961944 |  |  |
| 28 | 1950 | c.5849C>T | A1950V | De novo | - | - | 35982159 | Yes | 2 |

**Supplementary Table 2.**
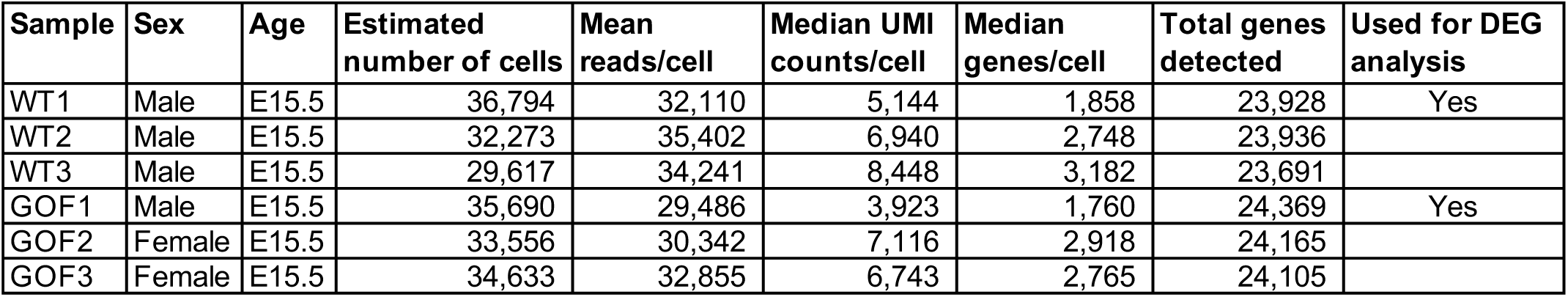
scRNA-seq summary.

**Supplementary Figure 1.**
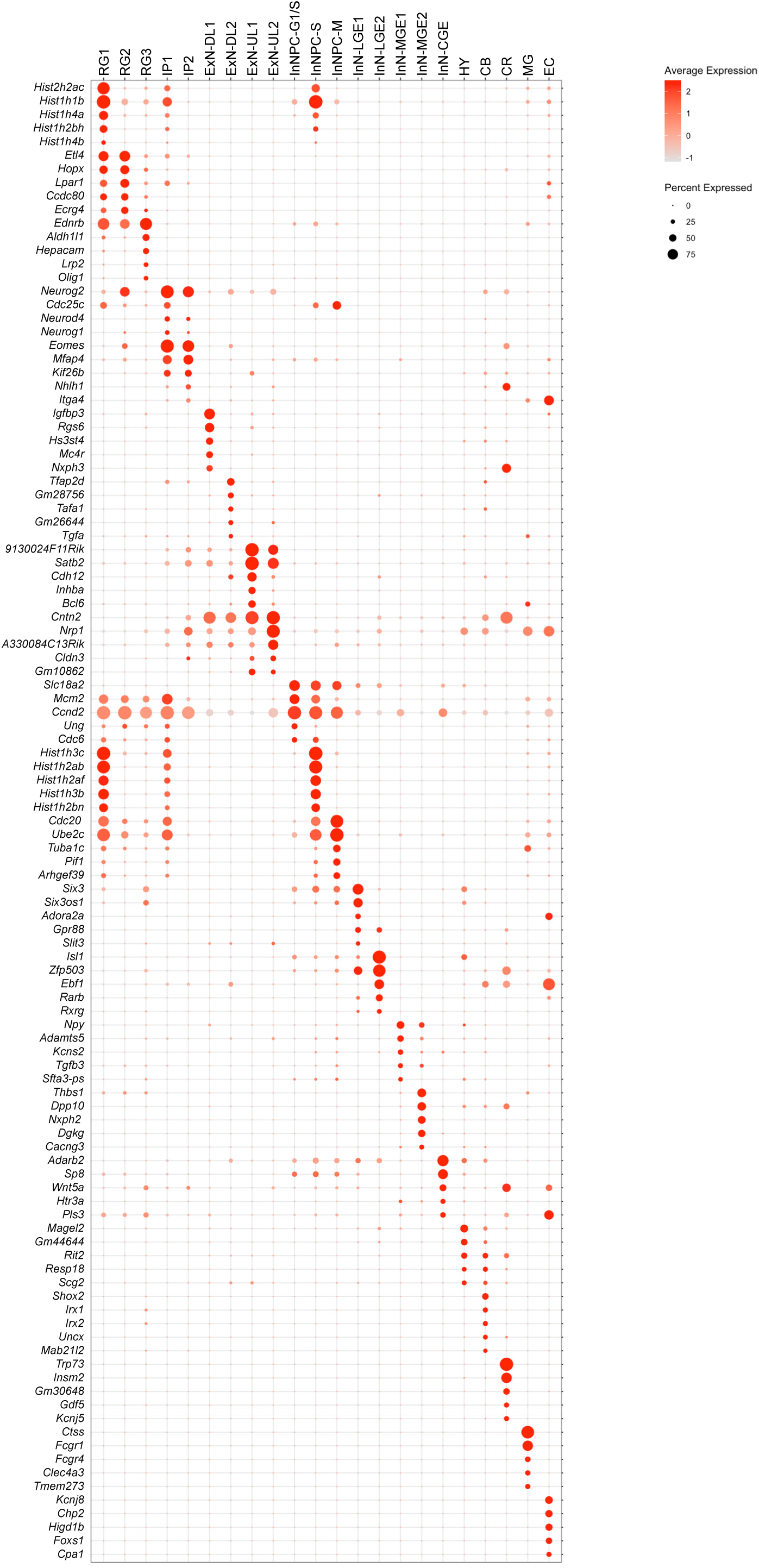
Five most highly expressed genes in each cell cluster.

## REFERENCES

1. Khachadourian, V., et al. Comorbidities in autism spectrum disorder and their etiologies. Transl Psychiatry 13, 71 (2023).

2. Courchesne, E., et al. Mapping early brain development in autism. Neuron 56, 399–413 (2007).

3. Courchesne, E., et al. Neuron number and size in prefrontal cortex of children with autism. JAMA 306, 2001–2010 (2011).

4. Hazlett, H.C., et al. Early brain development in infants at high risk for autism spectrum disorder. Nature 542, 348–351 (2017).

5. Courchesne, E., et al. The ASD Living Biology: from cell proliferation to clinical phenotype. Mol Psychiatry 24, 88–107 (2019).

6. Amaral, D.G., et al. In pursuit of neurophenotypes: The consequences of having autism and a big brain. Autism Res 10, 711–722 (2017).

7. Lainhart, J.E., et al. Head circumference and height in autism: a study by the Collaborative Program of Excellence in Autism. Am J Med Genet A 140, 2257–2274 (2006).

8. Courchesne, E., et al. Embryonic origin of two ASD subtypes of social symptom severity: the larger the brain cortical organoid size, the more severe the social symptoms. Mol Autism 15, 22 (2024).

9. Jourdon, A., et al. Modeling idiopathic autism in forebrain organoids reveals an imbalance of excitatory cortical neuron subtypes during early neurogenesis. Nat Neurosci 26, 1505–1515 (2023).

10. Chen, Y., Huang, W.C., Sejourne, J., Clipperton-Allen, A.E. & Page, D.T. Pten Mutations Alter Brain Growth Trajectory and Allocation of Cell Types through Elevated beta-Catenin Signaling. J Neurosci 35, 10252–10267 (2015).

11. Gompers, A.L., et al. Germline Chd8 haploinsufficiency alters brain development in mouse. Nat Neurosci 20, 1062–1073 (2017).

12. Bond, J., et al. ASPM is a major determinant of cerebral cortical size. Nature genetics 32, 316–320 (2002).

13. Jayaraman, D., Bae, B.I. & Walsh, C.A. The Genetics of Primary Microcephaly. Annu Rev Genomics Hum Genet 19, 177–200 (2018).

14. Kim, H.T., et al. The microcephaly gene aspm is involved in brain development in zebrafish. Biochem Biophys Res Commun 409, 640–644 (2011).

15. Jayaraman, D., et al. Microcephaly Proteins Wdr62 and Aspm Define a Mother Centriole Complex Regulating Centriole Biogenesis, Apical Complex, and Cell Fate. Neuron 92, 813–828 (2016).

16. Capecchi, M.R. & Pozner, A. ASPM regulates symmetric stem cell division by tuning Cyclin E ubiquitination. Nat Commun 6, 8763 (2015).

17. Pulvers, J.N., et al. Mutations in mouse Aspm (abnormal spindle-like microcephaly associated) cause not only microcephaly but also major defects in the germline. Proc Natl Acad Sci U S A 107, 16595–16600 (2010).

18. Fujimori, A., et al. Disruption of Aspm causes microcephaly with abnormal neuronal differentiation. Brain & development 36, 661–669 (2014).

19. Williams, S.E., et al. Aspm sustains postnatal cerebellar neurogenesis and medulloblastoma growth in mice. Development 142, 3921–3932 (2015).

20. Tonosaki, M., Fujimori, A., Yaoi, T. & Itoh, K. Loss of Aspm causes increased apoptosis of developing neural cells during mouse cerebral corticogenesis. PLoS One 18, e0294893 (2023).

21. Garrett, L., et al. A truncating Aspm allele leads to a complex cognitive phenotype and region-specific reductions in parvalbuminergic neurons. Transl Psychiatry 10, 66 (2020).

22. Johnson, M.B., et al. Aspm knockout ferret reveals an evolutionary mechanism governing cerebral cortical size. Nature 556, 370–375 (2018).

23. Kou, Z., et al. CRISPR/Cas9-mediated genome engineering of the ferret. Cell Res 25, 1372–1375 (2015).

24. Lindenhofer, D., et al. Cerebral organoids display dynamic clonal growth and tunable tissue replenishment. Nat Cell Biol 26, 710–718 (2024).

25. Li, R., et al. Recapitulating cortical development with organoid culture in vitro and modeling abnormal spindle-like (ASPM related primary) microcephaly disease. Protein Cell 8, 823–833 (2017).

26. van Benthem, A., et al. The microcephaly gene ASPM is required for the timely generation of human outer-radial glia progenitors by controlling mitotic spindle orientation. bioRxiv, 2023.2009.2025.559314 (2023).

27. Tsai, K.K., Bae, B.I., Hsu, C.C., Cheng, L.H. & Shaked, Y. Oncogenic ASPM Is a Regulatory Hub of Developmental and Stemness Signaling in Cancers. Cancer Res 83, 2993–3000 (2023).

28. Horvath, S., et al. Analysis of oncogenic signaling networks in glioblastoma identifies ASPM as a molecular target. Proc Natl Acad Sci U S A 103, 17402–17407 (2006).

29. Knol, M.J., et al. Genetic variants for head size share genes and pathways with cancer. Cell Rep Med 5, 101529 (2024).

30. Abrahams, B.S., et al. SFARI Gene 2.0: a community-driven knowledgebase for the autism spectrum disorders (ASDs). Mol Autism 4, 36 (2013).

31. Tate, J.G., et al. COSMIC: the Catalogue Of Somatic Mutations In Cancer. Nucleic Acids Res 47, D941–D947 (2019).

32. Krumm, N., et al. Excess of rare, inherited truncating mutations in autism. Nature genetics 47, 582–588 (2015).

33. Bae, B.I., Jayaraman, D. & Walsh, C.A. Genetic changes shaping the human brain. Dev Cell 32, 423–434 (2015).

34. McMillan, E.A., et al. Chemistry-First Approach for Nomination of Personalized Treatment in Lung Cancer. Cell 173, 864–878 e829 (2018).

35. Agam, Y., Joseph, R.M., Barton, J.J. & Manoach, D.S. Reduced cognitive control of response inhibition by the anterior cingulate cortex in autism spectrum disorders. NeuroImage 52, 336–347 (2010).

36. Haznedar, M.M., et al. Anterior cingulate gyrus volume and glucose metabolism in autistic disorder. Am J Psychiatry 154, 1047–1050 (1997).

37. Wang, X., Tsai, J.W., LaMonica, B. & Kriegstein, A.R. A new subtype of progenitor cell in the mouse embryonic neocortex. Nat Neurosci 14, 555–561 (2011).

38. Wang, L., Hou, S. & Han, Y.G. Hedgehog signaling promotes basal progenitor expansion and the growth and folding of the neocortex. Nat Neurosci 19, 888–896 (2016).

39. Zhao, X., et al. The transcriptional cofactor Tle3 reciprocally controls effector and central memory CD8(+) T cell fates. Nat Immunol 25, 294–306 (2024).

40. Ohtsuka, T. & Kageyama, R. Hes1 overexpression leads to expansion of embryonic neural stem cell pool and stem cell reservoir in the postnatal brain. Development 148 (2021).

41. Martinez-Cerdeno, V. Dendrite and spine modifications in autism and related neurodevelopmental disorders in patients and animal models. Dev Neurobiol 77, 393–404 (2017).

42. Fortier, A.V., Meisner, O.C., Nair, A.R. & Chang, S.W.C. Prefrontal circuits guiding social preference: Implications in autism spectrum disorder. Neurosci Biobehav Rev 141, 104803 (2022).

43. Levy, D.R., et al. Dynamics of social representation in the mouse prefrontal cortex. Nat Neurosci 22, 2013–2022 (2019).

44. Ahmed, S., et al. Transient impairment in microglial function causes sex-specific deficits in synaptic maturity and hippocampal function in mice exposed to early adversity. Brain Behav Immun 122, 95–109 (2024).

45. Buch, A.M., et al. Molecular and network-level mechanisms explaining individual differences in autism spectrum disorder. Nat Neurosci 26, 650–663 (2023).

46. Williams, Z.J., Suzman, E. & Woynaroski, T.G. Prevalence of Decreased Sound Tolerance (Hyperacusis) in Individuals With Autism Spectrum Disorder: A Meta-Analysis. Ear Hear 42, 1137–1150 (2021).

47. Truong, D.T., Rendall, A.R., Castelluccio, B.C., Eigsti, I.M. & Fitch, R.H. Auditory processing and morphological anomalies in medial geniculate nucleus of Cntnap2 mutant mice. Behav Neurosci 129, 731–743 (2015).

48. Platt, R.J., et al. Chd8 Mutation Leads to Autistic-like Behaviors and Impaired Striatal Circuits. Cell Rep 19, 335–350 (2017).

49. Yang, M., Lewis, F.C., Sarvi, M.S., Foley, G.M. & Crawley, J.N. 16p11.2 Deletion mice display cognitive deficits in touchscreen learning and novelty recognition tasks. Learn Mem 22, 622–632 (2015).

50. Peca, J., et al. Shank3 mutant mice display autistic-like behaviours and striatal dysfunction. Nature 472, 437–442 (2011).

51. Vasa, R.A. & Mazurek, M.O. An update on anxiety in youth with autism spectrum disorders. Curr Opin Psychiatry 28, 83–90 (2015).

52. Gerasimavicius, L., Livesey, B.J. & Marsh, J.A. Loss-of-function, gain-of-function and dominant-negative mutations have profoundly different effects on protein structure. Nat Commun 13, 3895 (2022).

53. Chen, S., et al. A genomic mutational constraint map using variation in 76,156 human genomes. Nature 625, 92–100 (2024).

54. Nicholas, A.K., et al. The molecular landscape of ASPM mutations in primary microcephaly. J Med Genet 46, 249–253 (2009).

55. Tunster, S.J. Genetic sex determination of mice by simplex PCR. Biol Sex Differ 8, 31 (2017).

56. Pachitariu, M., et al. Suite2p: beyond 10,000 neurons with standard two-photon microscopy. bioRxiv, 061507 (2017).

57. Pnevmatikakis, E.A., et al. Simultaneous Denoising, Deconvolution, and Demixing of Calcium Imaging Data. Neuron 89, 285–299 (2016).

58. Tomasi, D. & Volkow, N.D. Ultrafast method for mapping local functional connectivity hubs in the human brain. Annu Int Conf IEEE Eng Med Biol Soc 2010, 4274–4277 (2010).

59. Farina, M.G., et al. Small loci of astroglial glutamine synthetase deficiency in the postnatal brain cause epileptic seizures and impaired functional connectivity. Epilepsia 62, 2858–2870 (2021).

60. Sanganahalli, B.G., et al. Supraspinal Sensorimotor and Pain-Related Reorganization after a Hemicontusion Rat Cervical Spinal Cord Injury. J Neurotrauma 38, 3393–3405 (2021).

61. Kikusui, T., et al. Testosterone Increases the Emission of Ultrasonic Vocalizations With Different Acoustic Characteristics in Mice. Front Psychol 12, 680176 (2021).

